# MAVS is Important for Antiviral Defense Against Influenza A Virus in a Human Respiratory Epithelium Model

**DOI:** 10.64898/2025.12.09.693221

**Authors:** Maja Hemberg, Anne Louise Hansen, Jacob Storgaard, Julia Blay-Cadanet, Alice Pedersen, Anne Laugaard Thielke, Christian Kanstrup Holm

## Abstract

The respiratory epithelium is an important immunological barrier and the first line of defense against influenza A virus. In mice and in various cellular systems, induction of type I interferons (IFNα/β) during IAV infections is known to depend on cytosolic RNA sensors RIG-I and MDA5 and on the adaptor molecule MAVS. Until now, it has not been possible to directly test the importance of MAVS for induction of IFNs and for resistance to IAV infection in primary human respiratory epithelium.

Here, we used CRISPR-Cas9 to establish MAVS-deficient cultures of primary human respiratory epithelium using the air-liquid interphase culture system. Using this setup, we show that MAVS is indeed required for the induction of type I and type III IFNs and subsequently for the induction of IFN-stimulated genes in response to IAV infection in human primary respiratory epithelium. Finally, we demonstrate that MAVS is important for restricting viral replication in this model. In conclusion, we demonstrate that MAVS plays a non-redundant protective role during IAV infection in primary human respiratory epithelium.

## Introduction

The respiratory epithelium is the primary site of influenza A virus (IAV) infection and is a key initiator of antiviral defenses^1^. IAV is an important airborne virus, causing pandemics and seasonal endemics with great societal impact^2,3^. Vaccines and antivirals against IAV exist, however, vaccines must be renewed annually due to viral mutations, and antivirals are only effective if administered very early^2,4,5^. These challenges highlight the importance of understanding innate immunity, especially at the viral entry point, since this might provide insights to improve the development of more effective antiviral therapies.

Upon IAV infection, viral RNA is recognized by RIG-I-like receptors, including the retinoic acid-induced gene I (RIG-I) and melanoma differentiation-association gene 5 (MDA5)^6-9^. Both signal through the mitochondrial antiviral-signaling adaptor protein (MAVS, also known as ISP1, VISA or CARDIF^10-13^). Activation of MAVS leads to phosphorylation of the transcription factors interferon regulatory factors 3 and 7 (IRF3/7) and nuclear transcription factor-κB (NF-κB) that drive the production of interferons (IFNs) and subsequentially the induction of interferon stimulated genes (ISGs), which are essential for viral restriction^6,10^. MAVS is antagonized by IAV and other RNA viruses, emphasizing its importance in antiviral responses^14^. MAVS-deficiency has consistently demonstrated that MAVS is required for the production of type I IFNs (IFN- α and - β) during viral infections in cell types and mice^10,15-19^. However, there are discrepancies regarding the extent to which MAVS contributes to antiviral defense. Some studies show increased viral replication and higher susceptibility to RNA viruses in the absence of MAVS^10,15,17,20^, whereas others find no effect of MAVS KO on viral titers or survival following IAV infection^18,20^. Thus, the potential non-redundant protective role of MAVS during RNA virus infections remains unresolved. Only a few studies have examined the importance of MAVS in the antiviral defense against IAV, and these studies report no consistent phenotype^9,16,18,20^. Moreover, the function of MAVS in the respiratory epithelium, the first line of defense against IAV, has not been investigated. Understanding this is important for determining how early innate immune responses influence the outcome of IAV infection.

In this study, we demonstrate that a functional knockout of MAVS in a physiologically relevant model of human respiratory epithelium decreased production of type I and III IFNs and ISGs following IAV infection. Furthermore, we show that MAVS deficiency increases viral replication. These findings indicate that MAVS is important for the interferon response and protection against IAV infection in this human respiratory epithelium model.

## Results

### Functional KO of MAVS by CRISPR-Cas9 in HAE-ALI cultures

To investigate the importance of MAVS in respiratory epithelium we utilized a model where primary human airway epithelium cells (HAE) was cultured in the air-liquid interface (ALI) system. This model contains ciliated cells and mucus producing secretory cells which are hallmarks of respiratory epithelium, thus mimicking the biology of the human airway^21^. The model was established by collecting respiratory epithelial cells from the nasal cavity of donors and dedifferentiating them into basal cells. Genetic modification using CRISPR-Cas9 was then performed to generate MAVS knockout cells or AAVS1 knockout cells as a control. The KO cells were plated on Transwell membranes under liquid-liquid conditions. Once fully confluent, the apical medium was removed, initiating differentiation into a respiratory epithelium (Fig. 1A), containing ciliated and secretory cells (Fig. 1B). The KO was validated by Western blot, showing a decreased expression of MAVS in the MAVS KO culture compared to the AAVS1 KO culture (Fig. 1C). To test the functionality of the KO, we infected the KO HAE-ALI cultures with Sendai virus (SeV), which is known to trigger the RIG-I pathway^8,10^. Infection with SeV induced C-X-C Motif Chemokine Ligand 10 (CXCL-10) and Interferon Stimulated Gene 15 (ISG-15), measured by ELISA and qPCR, however this induction was decreased in the MAVS KO cultures compared to the control (Fig. 1D, E). The specificity of the KO was tested by stimulating the cultures with the STING ligand cGAMP, which is known to induce IFNs independently of the MAVS-pathway^22^. This resulted in a similar induction of CXCL-10 and ISG-15, measured by qPCR (Fig. 1F), and similar levels of Viperin, Interferon-Induced protein with Tetratricopeptide Repeats (IFIT1) and Interferon-Induced Transmembrane protein 3 (IFITM3) expression, measured by Western blot (Fig. 1G), between the KO cultures. This demonstrated that the KO of MAVS was specific and selectively affected the RIG-I and MDA-5 pathways in the HAE-ALI cultures. Next, we examined whether MAVS KO affected the cellular composition of the cultures. HAE-ALI cultures generated from two independent donors (donor 1 and donor 2) were assessed by qPCR for expression of Forkhead Box J1 (FOXJ1), Secretoglobin Family 1A Member 1 (SCGB1A1), Mucin 5AC (MUC5AC) and Mucin 5B (MUC5B) as markers of major respiratory epithelial cell types: ciliated cells, club cells, and goblet cells^23^. No differences were observed in the levels of ciliated or club cell markers between the KO cultures, however, the level of MUC5AC, a goblet cell marker, was lower in the MAVS KO culture from donor 2 compared to the AAVS1 KO. However, the level of MUC5B in this donor was similar between KO cultures, suggesting that the MUC5AC reduction was donor-specific and not a general effect of MAVS KO (Fig. 1H). To investigate cellular morphology, whole HAE-ALI membranes were fixed and stained with antibodies against protein markers of the various cell types. The membranes were imaged using fluorescence microscopy. No differences in morphology or composition were observed between AAVS1 or MAVS KO HAE-ALI cultures (Fig. 1I). These data suggest that MAVS-deficiency induced by CRISPR-Cas9 reduces the ISG response to SeV infection, indication that MAVS is important for the immune response in this model of human respiratory epithelium.

**Figure 1.**
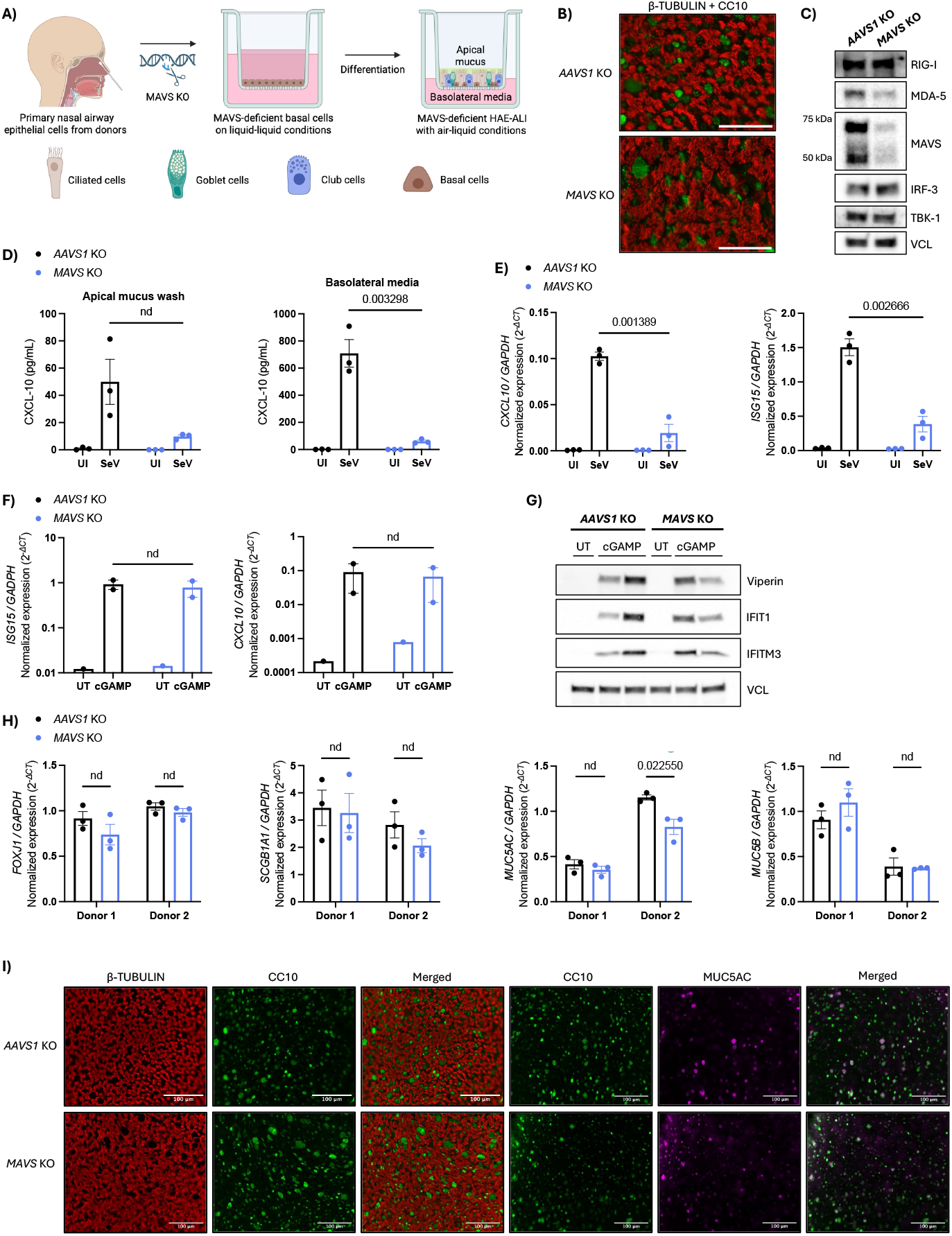
Functional knockout of MAVS by CRISPR-Cas9 in HAE-ALI cultures. **A)** Schematic of the human airway epithelium (HAE) cultured in an air-liquid interface (ALI) system. Epithelial cells were isolated from the nasal cavity of healthy donors. CRISPR-Cas9-mediated knockout of AAVS1 or MAVS was introduced in basal cells, which were subsequently seeded onto transwell membranes and differentiated into respiratory epithelium under ALI conditions. **B)** Immunofluorescence images of uninfected HAE-ALI showing cellular markers for ciliated cell in red (β-TUBULIN) and club cells in green (CC10). Scale bar = 50 µm. **C)** Western blotting of AAVS1 or MAVS KO HAE-ALI monolayer cultures using specific antibodies for the MAVS-pathway. Vinculin (VCL) was used as a loading control. **D)** AAVS1 KO and MAVS KO HAE-ALI cultures were infected with SeV (30 HAU/mL) or left uninfected for 16 hours. A hCXCL-10 ELISA was performed on the apical mucus wash and basolateral media. Each data point represent an independent culture (n=3) derived from the same donor. **E)** RT-qPCR analyzing ISG levels in the SeV infected or uninfected HAE-ALI. Data points represent independent cultures (n=3) from the same donor. **F, G)** AAVS1 or MAVS KO HAE-ALI cultures were stimulated with 2’3’-cGAMP (6 µg/mL) or left untreated for 16 hours. The cells were harvested and analyzed for ISGs using (E) RT-qPCR where each data point represents an independent culture (untreated n=1, treated n=2) or (F) Western blotting with VCL as loading control and with cell lysate from an independent culture derived from the same donor in each lane. **H)** RT-qPCR was performed on uninfected AAVS1 or MAVS KO HAE-ALI cultures derived from two independent donors for markers of ciliated cells (*FOXJ1*), club cells (*SCGB1A1*) or goblet cells (*MUC5AC* and *MUC5B*). Each data point represents one independent donor-derived culture (n=3). **I)** Immunofluorescence images of uninfected HAE-ALI showing cellular markers for ciliated cell in red (β-TUBULIN), club cells in green (CC10) or goblet cells in magenta (MUC5AC). Representative images are shown. Scale bar = 100 µm. For all panels, bars represent mean ± s.e.m. Statistical differences were determined by multiple unpaired t-tests with FDR correction using the two-stage step-up method of Benjamini, Krieger and Yekuteili (desired FDR = 5%). Q-values are shown above each comparison; “nd” indicates q-value > 0,05. Statistical analysis was not performed for untreated or uninfected groups (C-E).

### MAVS is essential for the interferon response in HAE-ALI after IAV infection

To test whether the absence of MAVS is important for the interferon response in respiratory epithelium during IAV infection, we infected the HAE-ALI cultures with IAV (A/PR/8/34, H1N1) for 16 hours or 48 hours. At 16 hours post-infection the induction of type I (IFN-β) and type III interferons (IFN-*λ*), measured by qPCR, was decreased in the MAVS KO cultures compared to the control (Fig. 2A). The same reduction was observed in the MAVS KO cultures at 48 hours post-infection and these findings was consistent across two separate donors (Fig. 2B). Furthermore, qPCR analysis of CXCL10 and ISG15 by qPCR showed that both genes were significantly reduced in the MAVS KO cultures compared to the control at 16 hours (Fig. 2C) and 48 hours (Fig. 2D) post-infection. When infecting with either IAV or SeV, Western blot analysis showed an induction of Viperin, IFITM3, ISG15 and IFIT1 in the AAVS1 KO cultures following 16 hours of infection. This induction was decreased in the MAVS KO cultures alongside the MAVS protein levels, but the induction of STING was similar between KO cultures (Fig. 2E). Reduced expression of ISGs was also observed at 48 hours post IAV-infection, although this reduction was less pronounced (Suppl. Fig. C). The release of CXCL10 from the HAE-cultures was examined by ELISA. This revealed a decreased amount of CXCL10 in the apical mucus wash as well as the basolateral media from the MAVS KO cultures compared to the control at both 16 hours (Fig. 2F, G) and at 48 hours post-IAV infection (Fig. 2H, I). Collectively, the lack of MAVS impaired the induction of IFNs and ISGs, demonstrating that the MAVS pathway is important for initiating the immune response against IAV in this model of respiratory epithelial cells.

**Figure 2.**
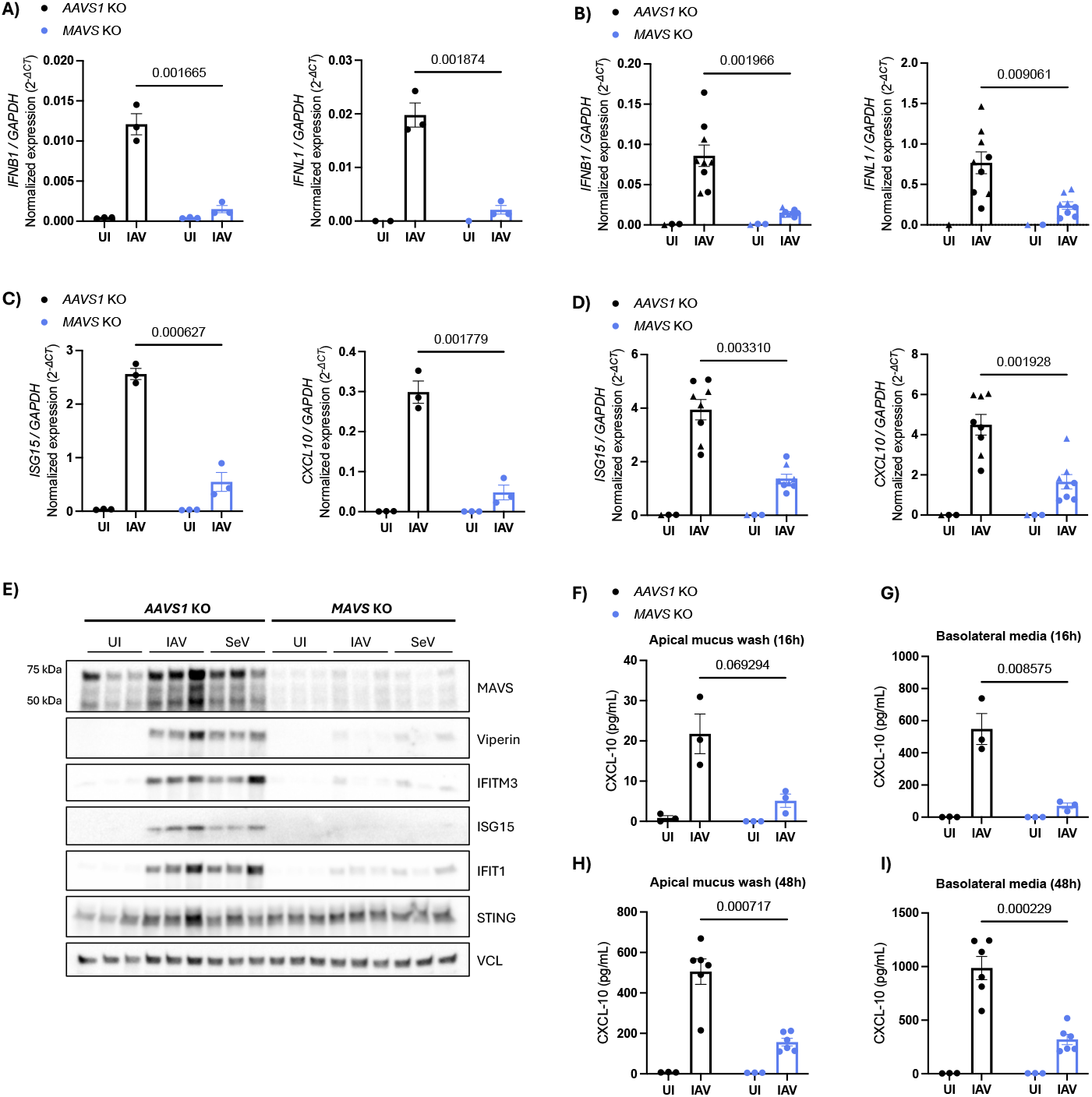
MAVS is essential for the interferon response in HAE-ALI after IAV infection. **A)** RT-qPCR analysis of *IFNB1* and *IFNL1* in AAVS1 or MAVS KO HAE-ALI cultures infected with IAV (diluted 1:20 in DMEM) or left uninfected for 16 hours. Each data point represents an independent culture (n=3) derived from the same donor. Data is representative of two independent experiments (see Suppl. Fig. A). For *IFNL1*, some uninfected cultures had undetectable levels. **B)** RT-qPCR analysis of *IFNB1* and *IFNL1* in AAVS1 or MAVS KO HAE-ALI cultures infected with IAV (1:20 in DMEM) or left uninfected for 48 hours. Pooled data from two independent experiments (donor 1 = • (uninfected n=2, infected n=4), donor 2 = ▴ (uninfected n=1, infected n=5)). Each data point represents an independent culture. Statistical significance was determined by multiple paired t-test (FDR=5%). **C)** RT-qPCR analysis on ISGs in cultures infected with IAV or left uninfected (like in A) for 16 hours. Each data point represents an independent culture (n=3) derived from the same donor. Data is representative of two independent experiments (see Suppl. Fig. B). **D)** RT-qPCR analysis of ISGs in AAVS1 or MAVS KO HAE-ALI cultures infected with IAV (1:20 in DMEM) or left uninfected for 48 hours. Pooled data from two independent experiments (donor 1 = • (uninfected n=2, infected n=4), donor 2 = ▴ (uninfected n=1, infected n=4)). Each data point represents an independent culture. Statistical significance was determined by multiple paired t-test (FDR=5%). **E)** Western blot analyzing protein expression of MAVS, various ISGs and STING in AAVS1 or MAVS KO HAE-ALI cultures infected with either IAV (diluted 1:20 in DMEM) or SeV (30 HAU/mL) or left uninfected for 16 hours. Vinculin (VCL) was used as a loading control. Each lane represents an independent culture (n=3) derived from the same donor. **F)** A hCXCL-10 ELISA was performed on cell-free apical mucus wash from AAVS1 or MAVS KO HAE-ALI cultures infected with IAV (1:20 in DMEM) or left uninfected for 16 hours. Each point represents an independent culture (n=3) from the same donor. **G)** Basolateral media from the same experiment as (F) analyzed by hCXCL-10 ELISA (n=3). **H)** hCXCL-10 ELISA on cell-free apical mucus wash from AAVS1 or MAVS KO HAE-ALI cultures infected with IAV (1:20 in DMEM) or left uninfected for 48 hours. Each data point represents an independent culture (uninfected n=3, infected n=6) from the same donor. Data representative of two independent experiments (see Suppl. Fig. D). **I)** hCXCL-10 ELISA on basolateral media from the same experiment as (H). Each data point represents an independent culture (uninfected n=3, infected n=6) derived from the same donor. Data representative of two independent experiments (see Suppl. Fig. E). For all panels, bars represent mean ± s.e.m. Unless otherwise stated, statistical differences were determined by multiple unpaired t-tests with FDR correction using the two-stage step-up method of Benjamini, Krieger and Yekuteili (desired FDR = 5%). Q-values are shown above each comparison; “nd” indicates q-value > 0,05. Statistical analysis was not performed for uninfected groups (A-D and F-I).

### MAVS is non-redundant for the protection against IAV in HAE-ALI

Considering that MAVS was important for the IFN response in the HAE-ALI, we wanted to investigate if the reduced MAVS expression also affected the viral replication in these cultures. To examine this, viral RNA corresponding to the NP, M2, and NS1 segments of IAV from infected AAVS1 or MAVS KO HAE-ALI was quantified by qPCR. At 48 hours post-IAV infection, viral RNA levels were increased in the MAVS KO cultures compared to the control for all viral genes. This finding was consistent across two independent donors (Fig. 3A). This effect of MAVS KO was observed only after longer infections, as viral gene levels were similar between the two KO cultures at 16 hours post-infection (Suppl. Fig. F). The expression of viral NS1-protein was analyzed by Western blot, showing an increase of NS1 in the MAVS KO culture compared to the control (Fig. 3B). This finding was consistent across two separate experiments and quantification showed a significant increase in the NS1 expression normalized to the loading control (Fig. 3C). Next, we examined the entire HAE-ALI membrane by immunofluorescence. Infected or uninfected AAVS1 or MAVS KO cultures were fixed 16 hours after infection or mock infection. The membranes were stained with DAPI and an antibody against the IAV protein NS1, and imaged using a fluorescence microscope. Quantification of NS1-positive cells showed that approximately 10% of cells were NS1-positive in AAVS1 KO cultures, whereas MAVS KO cultures exhibited a higher mean of approximately 13% NS1-positive cells (fig. 3D, E). Based on the immunofluorescence images, it was not possible to determine if any specific cell type was more likely to be infected, however, viral protein was observed in cells with a ciliated appearance in both cultures (Fig. 3F). Together, increased viral RNA levels, higher accumulation of viral protein by Western blot, and a greater proportion of NS1-positive cells indicate that the viral replication of IAV is enhanced in this MAVS-deficient respiratory epithelium model.

**Figure 3.**
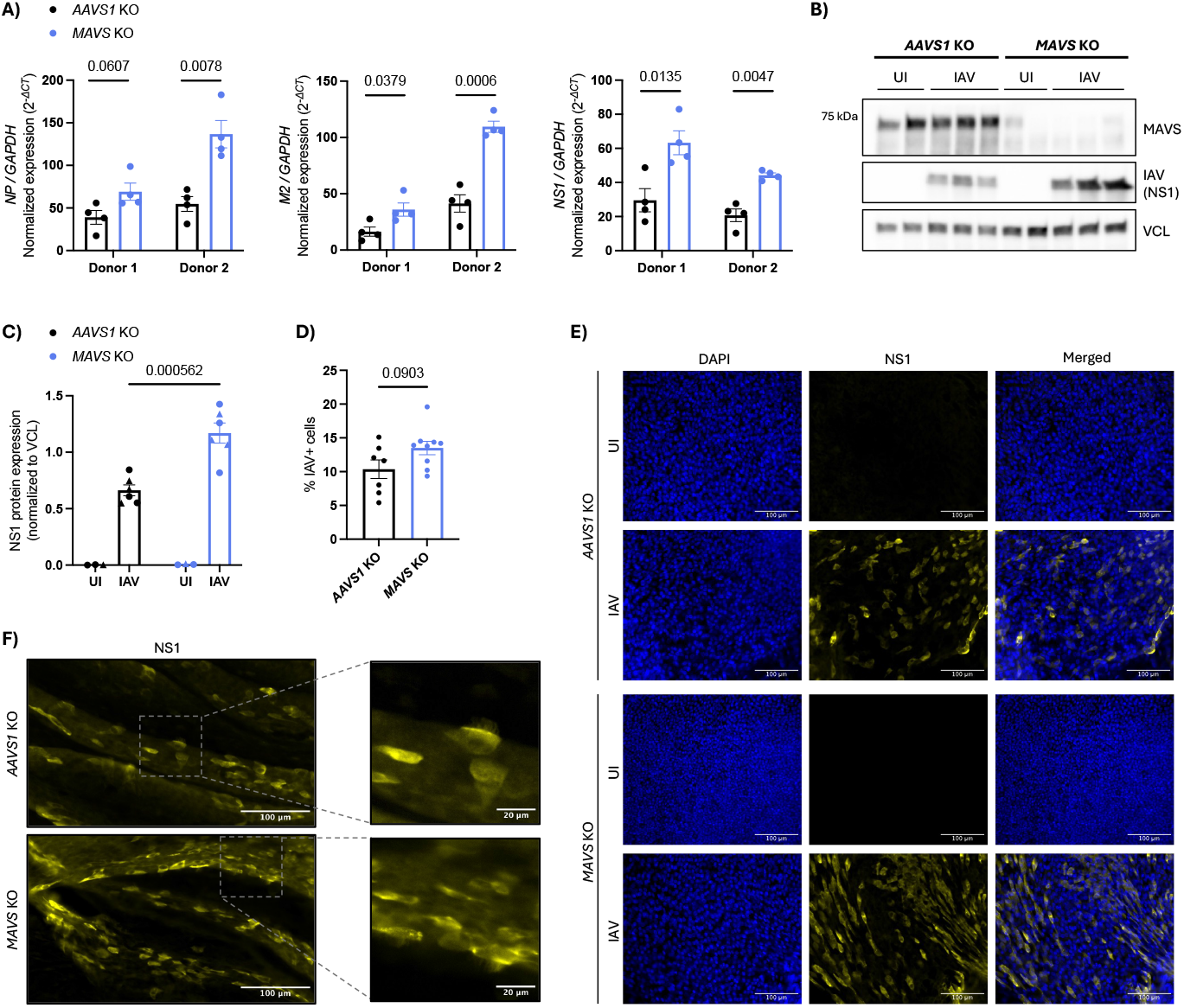
MAVS is non-redundant for the protection against IAV in HAE-ALI. **A)** RT-qPCR analysis of viral RNA from three different IAV segments in AAVS1 or MAVS KO HAE-ALI cultures infected with IAV (diluted 1:20 in DMEM) for 48 hours. Each data point represents an independent culture (n=4) derived from two different donors. Statistical significance was determined using a Welch’s t-test. **B)** Western blot analysis of viral (NS1) and MAVS protein levels in AAVS1 or MAVS KO HAE-ALI cultures infected with IAV (1:20 in DMEM) or left uninfected for 48 hours. Vinculin (VCL) was used as a loading control. Each lane represents an independent culture (uninfected n=2, infected n=3) derived from the same donor. Data is representative of two independent experiments (see suppl. fig. C). **C)** Quantification of the NS1 protein levels determined by western blotting and normalized to the loading control (VCL). Pooled data from two independent experiments (donor 1 = • (uninfected n=2, infected n=3), donor 2 =▴ (uninfected n=1, infected n=3)). Each data point represents an independent culture. Statistical significance was determined using a multiple unpaired t-test (FDR=5%). **D)** Quantification of immunofluorescence (IF) images of AAVS1 or MAVS KO HAE-ALI cultures 16 h post-infection with IAV (1:20 in DMEM). The graph shows the proportion of NS1+ cells relative to the total cell count. Cell counting was performed in ImageJ. Each data point represents one 40x IF image (AAVS1 KO, n=12; MAVS KO, n=13) from a single culture per condition. Statistical difference was determined using a Welch’s t-test, and the p-value is shown above the comparison. **E)** Immunofluorescence images acquired at 40x magnification of AAVS1 or MAVS KO HAE-ALI cultures derived from the same donor infected with IAV or left uninfected for 16 hours. Nuclei were stained with DAPI (blue) and viral NS1 protein with an anti-NS1 antibody (yellow). Representative images of each condition are shown. Scale bar = 100 µm. **F)** IF images acquired at 40x magnification of AAVS1 or MAVS KO HAE-ALI cultures 16 h post-infection with IAV. Images are derived from the same donor. A magnified inset of boxed of regions are shown to the right. Scale bars 100 µm (main) and 20 µm (insets). Bars indicate mean ± s.e.m. FDR correction by the two-stage step-up method of Benjamini, Krieger and Yekuteili (desired FDR = 5%). The q-values are indicated above comparisons for (A) and (C). Statistical analysis was not performed for uninfected groups (C).

## Discussion

Together, our results demonstrate establishment of a physiological relevant MAVS-deficient model of the respiratory epithelium. The specific role of MAVS in respiratory epithelium has not previously been reported, but here we show that MAVS is important for the production of IFNs and ISGs following IAV infection in this respiratory epithelium model. Furthermore, we show that viral replication was increased during prolonged infection in the absence of MAVS.

Consistent with our results, most studies examining the role of MAVS during infection with RNA viruses, show a decreased induction of type I and type III interferons following MAVS KO or silencing of MAVS^10,15-19^. Many studies also report an effect of the MAVS-deficiency on the production of ISGs. Regarding the antiviral properties of MAVS, some studies indicate that MAVS is important for protection against various RNA viruses^10,15,17^. The importance of MAVS in IAV infections has only been investigated in few studies, showing that MAVS is essential for the induction of IFNs and ISGs, supporting the findings of the current study. However, most studies report no observable effect of MAVS KO on antiviral protection^18,20^. Those studies were conducted in MAVS KO mice, so it remains unclear whether the findings apply to primary human cells. Moreover, they do not allow conclusions about the isolated role of MAVS in the respiratory epithelium for the protection against IAV.

To our knowledge, this is the first study examining the importance of MAVS in a model of the human respiratory epithelium. Our findings suggest that in the absence of MAVS, viral RNA sensing by RIG-I or MDA5 fails to trigger the induction of IFNs and ISGs. This induction, following recognition of viral RNA, appears to be important for restricting viral replication in the respiratory epithelium during prolonged IAV infections. The smaller induction of IFNs and ISGs that does occur in the MAVS KO cultures might be a result of other signaling pathways or some HAE-ALI cells that wasn’t affected by the KO.

The HAE-ALI cultures are heterogeneous, and electroporation may not affect all cells, making it difficult to generate a fully MAVS-deficient culture. Nonetheless, MAVS protein levels were reduced in all KO cultures for both donors compared to the control. This reduction allowed us to investigate the importance of MAVS deficiency in this respiratory epithelium model. To examine possible off-target effects of the KO, we have presented that MAVS KO did not affect the cellular composition of the model. The current study only included two same-sex donors. More donor variability could have strengthened the results even further. The HAE-ALI model includes only respiratory epithelial cells, and therefore it does not capture the contribution of immune cells or the interplay between epithelium and immune cells in antiviral protection. However, based on this study, we can estimate the isolated role of MAVS in respiratory epithelium. Further research is needed to establish the importance of MAVS when including the immune cells. An important strength of this study is, that it was conducted on primary human respiratory epithelium cells, making the results transferable to humans.

Improving our understanding of the innate immunity, especially at the viral entry point, is important to aid the future development of antivirals against IAV. Here, the HAE-ALI culture has a lot of potential, since it allows knockout of individual protein in the innate immune response. Furthermore, this model uses primary human cells, limiting the use of animal experiments, because we can perform infectious studies in a model with complex humane tissue. In this study, we used this model to establish a non-redundant protective role for MAVS in human respiratory epithelium infected with IAV.

## Methods and materials

### Establishment of the air-liquid interface epithelium model

Primary nasal epithelial cells were isolated from healthy donors using an interdental brush (Brage Nilsson, P-My x-small). The brush was inserted in an angle following the bridge of the nose and twisted at the nasal turbinates. Cells were rinsed off the brush by gently expelling Monolayer medium (Airway Epithelial Cell Basal Medium (PromoCell, #C-21260) + 1 pack of Airway Epithelial Cell Growth Medium Supplement (PromoCell, #C-39160) + 100 U/mL Penicillin/Streptomycin (Gibco, #15140122)) and PBS (Biowest, #L0615). The cells were spun down at 130 x g for 5 min and resuspended in 2 mL Monolayer medium before being moved into a T10 flask (Thermo Scientific, #156367) precoated 24 hours earlier with 0,1 mg/mL Bovine type I collagen solution (Sigma-Aldrich, #804592, diluted in sterile ddH_2_O). Once the monolayer cells were approx. 80% confluent, they were split into a T75 flask (Sarstedt, #83.8311) pre-coated with collagen using 1x Trypsin with 0,3 mM EDTA (10x Trypsin, (Gibco, 2,5%, #15090), diluted to working conc. in PBS + UltraPure 0,5 mM EDTA (Invitrogen, #15575)). When the cells were confluent in the T75 flasks, they were ready for genetic alteration. The collection and culturing of primary cells were approved by the Danish Research Ethics Committees (Case nr.: 1-10-72-182-19). Both donors in this study were male.

### CRISPR-Cas9 KO

Monolayer cells were genetically modified twice to ensure efficient MAVS knockout, using two sequential rounds of CRISPR-Cas9-mediated knockout of either MAVS or AAVS1 (control). Both knockout rounds followed the same procedure; however, after the first KO, the cells were seeded into T75 flasks pre-coated with 0,1 mg/mL Bovine type I collagen solution and maintained in Monolayer medium until they had recovered and reached sufficient confluency for the second KO. Following the second KO, cells were seeded onto 6,5 mm Transwell membranes (Corning, #3470) pre-coated with 30 µg/mL Bovine type I collagen solution (diluted in sterile ddH2O) with 75,000 cells pr. membrane.

The monolayer cells were detached from their flasks using Trypsin/EDTA. Trypsin was inactivated by DMEM (low glucose, Gibco, #11880-028) with 5% FBS. Cells were spun down at 130 x g for 5 minutes, resuspended in Monolayer medium, and counted using a Bürker-Türk chamber. Genetic modification was done by CRISPR-Cas9 KO of MAVS and AAVS1 (control). The sgRNAs used for the experiments were purchased from Synthego:

AAVS1 sgRNA: 5’-GGGGCCACUAGGGACAGGAU-3’

MAVS sgRNA #1: 5’-UGUCUUCCAGGAUCGACUGC −3’

MAVS sgRNA #2: 5’-CCGGUUCCCUGAGAGUGUGC −3’

The sgRNAs were resuspended and their concentrations were adjusted to 3200 ng/µL using TE-buffer (provided with sgRNAs). RNP complexes for each sgRNA were prepared in PCR tubes by mixing 0.6 µL of Alt-R S.p. Cas9 nuclease V3 (IDT, # 1081059) with 3,2 µg of sgRNA pr. electroporation (max. 200.000 cells pr. electroporation). After 15 minutes at room temperature, the RNP complexes for MAVS #1 and MAVS #2 were mixed. For the initial KO, the entire cell suspension was used, however, for the second KO, a volume of the cell suspension corresponding to 75.000 cells pr. membrane was used. The cells were spun at 130 x g for 5 minutes, resuspended with PBS and spun down once more. After removing the supernatant, the cells were resuspended in 20 µL Opti-MEM (Thermo Fisher, #31985062) pr. electroporation. The cell suspension in Opti-MEM was divided between the two PCR tubes containing the AAVS1 or MAVS RNP-complexes.

The solutions were added to an electroporation strip (Lonza, Nucleocuvette Strips, #V4XC-2032) and electroporated using the Lonza 4D-Nucleofector with the program DC.100 and the cell type P3. After electroporation, the cells rested for 2 minutes. Monolayer medium or Submerged medium (DMEM + 200 U/mL Pen/Strep mixed 1:1 with 2x ALI medium (Airway Epithelial Cell Basal Medium (PromoCell, #C-21260) + 2 packs of Airway Epithelial Cell Growth Medium Supplement (PromoCell, #C-39160) but without triiodothyronine + 1 mL of 1.5 mg/mL BSA) was used to remove the cells from the electroporation strip. The cell suspension in Monolayer medium (1^st^ KO) was added two separate T75 flasks. The cell suspension from the 2^nd^ KO was seeded and submerged with Submerged medium on Transwell membranes. When cultures reached full confluency, ALI (Ali-Liquid Interface) was introduced and ALI medium (Pneumacult ALI medium kit (StemCell, #05002) + ALI medium supplement (StemCell, #05003) + 100 U/mL Pen/Strep, supplemented with 24 µg of hydrocortisone (StemCell, #07925) and 0.2 mg heparin (StemCell, #07980)) was added to basolateral compartment. The cells differentiated for at least 21 days, verified by ciliated beating and mucus production. Cells were kept at 37°C, 5% CO_2_. Throughout the establishment and KO of the model, media was changed three times a week. Once movement of the cilia was observed, the cultured were washed weekly with DMEM to remove excess mucus.

### Infection or treatment of HAE-ALI

Before treatment or infection, the HAE-ALI cultures were washed for 5 minutes with DMEM to remove mucus build-up, and the ALI medium in the basolateral compartment was changed. For cGAMP treatment, 150 µl of DMEM or a dilution of 6 µg/mL 2’3’-cGAMP (InvivoGen, #tlrl-nacga23-02) in DMEM was added to the apical surface. After 1 hour in the incubator at 37°C, the apical solutions were removed, and the cultures were placed in the incubator for 16 hours. Infection of the HAE-ALI was performed by adding a virus dilution in DMEM, or by mock-infecting the cultures with DMEM, for 1 hour at 37°C before removing the solutions a returning the cultures in the incubator for 16 or 48 hours.

At time of harvest, the HAE-ALI cultures were washed for 5 minutes with DMEM and the basolateral media was removed, saving both the wash-solutions and media for ELISA. 400 µL of Trypsin/EDTA was added basolaterally and 150 µL was added to the apical surface. After approximately 5 minutes, the cells were resuspended in the Trypsin/EDTA solution and transferred to a tube with 500 µL of DMEM with 5% FBS. Cells that remained attached to the membrane were gently scraped off with the pipette tip. Cells were pelleted at 300 x g for 5 minutes, resuspended in cold PBS, and divided into two tubes. The tubes were centrifuged again at 300 x g for 5 minutes, after which the cells were lysed for either RNA isolation or Western blot.

### Viruses

For the infection studies, we used Sendai virus, Cantell strain (#10100816, Charles River) with 6000 HAU/ml, which was diluted 1:200 in DMEM (30 HAU/ml). Furthermore, we used the Influenza A /PR/8/34 (H1N1) strain (#10100374, Charles River) with an MOI of 0.5.

### Westen Blotting

We used Western Blotting to detect proteins in our samples. Protein was isolated from the samples using a lysis buffer consisting of ice-cold RIPA lysis buffer (Thermo Fisher Scientific, #89901) supplemented with 5 IU/mL Benzonase (Sigma-Aldrich, #E1014), 10 mM Sodium Fluoride, and 1x complete protease inhibitor cocktail (Roche, #5892988001). Protein concentrations were determined using a BCA protein assay kit (Thermo, #23225). Before Western Blotting, the sample was mixed with XT Sample Buffer (Bio-Rad, #1610791) and XT Reducing Agent (Bio-Rad, #161-0792) and denatured at 95°C for 2-3 minutes. The amount of protein loaded in each well was approximately 7,5 µg and the Precision Plus Protein Dual Color Standard (Bio-Rad, #161-0374) was used as a protein ladder. Using a 4–20% Criterion TGX Precast Midi Protein Gel (Bio-Rad, #5671094 or #5671095), samples was separated by SDS-Page running at 70V for approx. 15 min followed by approx. 50-70 min at 120V. The protein was transferred to a PVDF membrane (Bio-Rad, #170-4157) using a Trans-Blot Turbo Transfer from Bio-Rad. The membranes were blocked for 1 hour at room temperature using PBS/Tween (PBS, Biowest, #X0515, added to 4,5 L of deionized H_2_O) with 0.05% Tween 20 (Sigma-Aldrich, P1379) (PBS/Tween) mixed with 5% skim-milk powder (Sigma Aldrich, #70166). After blocking, the membranes were rinsed with PBS/Tween and cut to match the molecular weight range of the protein of interest. The membrane pieces were incubated ON at 4°C with the following antibody solutions diluted in PBS/Tween: anti-RIGI (Cell Signaling, #3743, 1:1000), anti-MDA5 (Cell Signaling, #5321, 1:1000), anti-MAVS (Cell Signaling, #24930, 1:1000), anti-IRF3 (Cell Signaling, #11904, 1:1000), anti-TBK1 (Cell Signaling, #3013S, 1:1000), anti-Viperin (Cell Signaling, #13996, 1:1000), anti-IFIT1 (Cell Signaling, #14769, 1:1000), anti-IFITM3 (Cell Signaling, #59212, 1:1000), anti-ISG15 (Cell Signaling, #2758, 1:1000), anti-NS1 (Invitrogen, #PA5-32243, 1:1000), or anti-STING (Cell Signaling, #13647). Anti-Vinculin (Sigma, #V9131, 1:10000) was used as a loading control. After three washes with PBS/Tween, the membranes were incubated with secondary antibody solutions with peroxidase-conjugated F(ab)2 donkey anti-mouse IgG (Jackson, #711-036-150, 1:10000) or peroxidase-conjugated F(ab)2 donkey anti-rabbit IgG (Jackson, #711-036-152, 1:10000) for 1-2 hours at room temperature. The membranes were images using an Image Quant LAS4000 mini (GE Healthcare), exposing which either SuperSignal West Pico PLUS chemiluminescent substrate (Thermo, # 34577) or the SuperSignal West Femto chemiluminescent substrate (Thermo, #34096).

### Reverse transcriptase-quantitative polymerase chain reaction (RT-qPCR)

RT-qPCR was used to quantify the transcription of genes of interest. RNA was lysed and isolated from the samples using the High Pure RNA Isolation Kit (Life Science, #11828665001) according to the manufacturer’s protocol. RNA concentrations were measured with a NanoDrop spectrophotometer (Thermo Fisher). Gene expression was analyzed using premade TaqMan assays and the RNA-to-Ct-1-Step kit (Applied Biosciences), following manufacturer’s recommendations. The RT-qPCR was performed on an AriaMx Real-Time PCR System. TaqMan assays for qPCR were purchased from Applied Bioscience; GAPDH (Hs02786624), CXCL10 (Hs00171042), ISG15 (Hs01921425), FOXJ1 (Hs00230964), SCGB1A1 (Hs00171092), MUC5AC (Hs01365616), MUC5B (Hs06629268) IFNB1 (Hs01077958), and IFNL1 (Hs00601677). Furthermore, we used custom made TaqMan primers for:

NP – segment 5: Forward 5’-GGAAATTTCAAACTGCTGCACAAAA, Reverse 5’- CGTGCTAGAAAAGTGAGATCTTCGA

M2 – segment 7: Forward 5’-AACCTGTGAACAGATTGCTGACT, Reverse 5’- TCTGATTAGTGGGTTGGTTGTTGT

NS1 – segment 8: Forward 5’- GCTAAGGGCTTTCACCGAAGAG, Reverse 5’- TGGAAGAGAAGGCAATGGTGAAATT

### hCXCL-10 ELISA

To quantify the release of CXCL-10 from the HAE-ALI cultures, we used hCXCL-10 ELISA. The ELISAs were performed on apical mucus washes and on basolateral media collected prior to cell harvest. A 96 well, half-area, flat-bottom plate (Corning, 3690, VWR) was coated with a capture antibody mix from the CXCL10 DuoSet ELISA (R&D, #DY266), prepared according to manufacturer’s protocol, and incubated ON at room temperature. The plate was washed with a wash buffer consisting of PBS (Biowest, #X0515, mixed with 4,5 L of deionized H_2_O) with 0.05% Tween 20 (Sigma-Aldrich, P1379). This wash was used three times between each step of the ELISA. The plate was blocked for a minimum of 1 hour using a Reagent Diluent with 1% BSA (Sigma, #9048-46-8) in PBS. The standard dilutions (R&D, #DY266, prepared using manufacturer’s protocol), blanks and sample dilutions (undiluted for 16 hours, diluted 1:10 for mucus 48 h p.i. and 1:20 for media 48 h p.i.) was added and incubated ON at 4°C. The following day, a detection antibody mix (R&D, #DY266, prepared using manufacturer’s protocol) was added and left to incubate at room temperature for 2 hours. Next, a 1:40 dilution of Streptavidin-HPR B (R&D, #DY266, Part-number 893975) in Reagent Diluent was added for 20 min at RT in the dark. After a final wash, 50 µL of the substrate solution TMB One Solution (Promega, #G7431) was added and incubated in the dark until the color development was sufficient. The reaction was stopped after approx. 10-20 minutes by adding 25 µl of stop-solution (1 mol/L (2N) sulfuric acid (H_2_SO_4_), VWR Chemicals). Results were acquired with a plate reader (BioTek Synergy HTX Multimode Reader) with optical density determined at 450 nm, using 570 nm for wavelength correction.

### Fluorescence staining of HAE-ALI cultures

Before staining, the AAVS1 KO and MAVS KO HAE-ALI membranes were washed for 5 minutes to remove mucus build-up by adding 200 µL of DMEM. The basolateral media was removed and the Transwell inserts were moved into a new 24-well plate (Sarstedt, #83.3922). Cells were fixed using 4% formaldehyde solution (16 % formaldehyde solution, Methanol-free, (Pierce #28908), diluted in PBS) by adding 200 µL of the solution to the apical surface and 400 µL to the basolateral compartment for 20-30 minutes at room temperature. After fixation, both compartments were washed three times with PBS. Plates can be stored at 4°C for approx. 1 month if 250 µL PBS with Sodium azide (1:100 dilution) is added to the apical surface and 800 µL to the basolateral compartment. Before staining, the cells were permeabilized by adding freshly made 0.2 % Triton-X100 (in PBS) to the apical surface and blocked using 0,5% BSA (in PBS). After blocking, the membranes were removed from the Transwell inserts by cutting around the outer edge with a scalpel. Each membrane was divided into four pieces, that were stained separately. Primary antibody solutions (diluted in the 0,5% BSA blocking buffer) were added for 1-2 hours. After three washes with blocking buffer, the membrane pieces were incubated for 1 hour in the dark with secondary antibodies diluted 1:500 in blocking buffer. For some of the membranes, a DAPI stain was added during the last 5 minutes of the incubation. The staining of the membrane pieces was as follows:

- Primary: Anti-β-tubulin (Abcam, #Ab179509, 1:500) and anti-CC10 (Santa Cruz Biotechnology, #sc-365992, 1:200). Secondary: Anti-rabbit-Alexa Flour 488 (Invitrogen, #A-32790) and anti-mouse Alexa Fluor 647(Invitrogen, #A-21463).
- Primary: Anti-MUC5AC (Cell Signaling, #61193, 1:200) and anti-CC10 (Santa Cruz Biotechnology, #sc-365992, 1:200). Secondary: Anti-rabbit-Alexa Flour 488 (Invitrogen, #A-32790) and anti-mouse Alexa Fluor 647(Invitrogen, #A-21463).
- Primary: Anti-HA (Abcam, #ab8262, 1:500). Secondary: Anti-mouse Alexa Fluor 647(Invitrogen, #A-21463) and DAPI (Sigma Aldrich, #D9542, 1:500)
- Primary: Anti-NS1 (Invitrogen, #PA5-32243, 1:200). Secondary: Anti-rabbit Alexa Fluor 647(Invitrogen, #A-21245) and DAPI Sigma Aldrich, #D9542, 1:500).

Before mounting, each membrane piece was washed three times with blocking buffer. One drop of ProLong Glas Antifade Mountant (Invitrogen, #P36982) was added to a glass slide. The membrane pieces were blotted against paper tissue and placed in the mounting media cell side up. A coverslip was added on top, and the slide was allowed to dry horizontally ON before imaging. Fluorescence images were acquired using an Olympus BX63 Upright Widefield Fluorescence microscope with a 40x, Plan Flurite objective and a Sensitive Andor Zyla 5.5 camera. Images were acquired and analyzed at the Bioimaging Core Facility, Health, Aarhus University, Denmark. All images were acquired using identical microscope settings.

Brightness and contrast were adjusted in ImageJ for visualization.

### Statistical analysis

Graphs and statistics were created and calculated using Graph Pad Prism 10. For all panels, bars represent mean ± s.e.m. Unless otherwise stated in the figure legend, statistical significance was assessed using multiple unpaired t-tests. The false discovery rate (FDR) in both unpaired and paired multiple t-tests was controlled by the two-stage step-up method of Benjamini, Krieger and Yekutieli with a desired FDR = 5%. Q-values are indicated above each comparison. Results with a q < 0.05 were considered significant.

### Quantification of Western Blotting

Membrane images from Western Blotting were imported into the AzureSpot Pro 1.5 software. Membranes blotted for either NS1 or Vinculin were analyzed separately. Lanes were defined by adding a box with the correct number of lanes and adjusting each box manually to include the band. Background was subtracted using “Rolling Ball” with a radius of 40. In all lane, the band was detected manually with a fixed band size of 30 pixels. The volumes of each band were exported to an Excel sheet where the volume of NS1 was normalized to the volume of Vinculin.

### Quantification of immunofluorescence images (IF)

DAPI- and NS1-stained cells were quantified from IF images acquired at 40x magnification. The images were imported into ImageJ2 (version 2.16.0), and macros were created to quantify cells numbers. The following ImageJ tools were applied to all DAPI-stained images: Subtract Background (rolling=100), Gaussian Blur (sigma=4 slice), Enhance Contrast (saturated=0.50 normalize), Auto Threshold (Default, dark), Convert to Mask, Watershed, Fill Holes, and Analyze Particles (size=10-Infinity). To estimate the number of NS1-positive cells, a second macro was applied: Subtract Background (rolling=100), Gaussian Blur (sigma=2 slice), Enhance Contrast (saturated=0.35 normalize), Auto Threshold (Triangle, dark), Convert to Mask, Close, Fill Holes, Distance Map, Find Maxima (prominence=10 exclude, output=Maxima Within Tolerance), and Analyze Particles (size=10-Infinity). For each macro, the “Count” value from the summary of each image was imported to an Excel sheet, and the number of NS1-positive cells was normalized to the total cell count.

## Acknowledgements

This work was supported by Frimodt-Heineke Fonden, Dagmar Marshalls Fond, the Independent Research Fond Denmark (4287-00003B) and Hallas Møller Ascending Investigator (#0066798). We would like to acknowledge the Bioimaging Core Facility, Health, Aarhus University, Denmark, for the use of equipment and support.

## Author contributions

C.H. designed the project. M.H planned and performed experiments, analyzed data, prepared figures and drafted the manuscript. A.L.H., J.S., J.B-C., A.P., and A.T. planned experiments and analyzed data. C.H. and M.H. finalized the manuscript.

## Supplementary Figures

**Figure S1.**
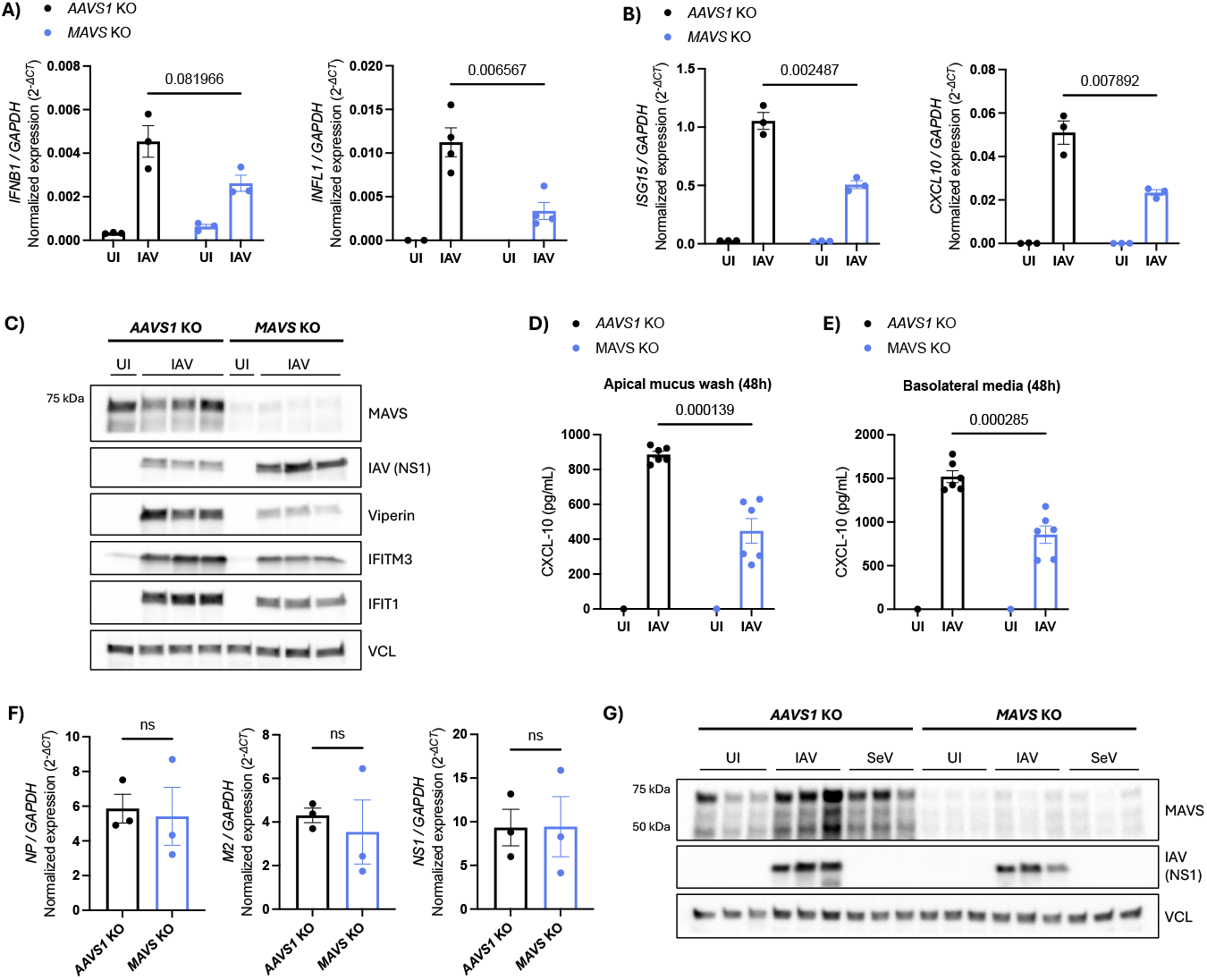
Supplementary data. **A)** RT-qPCR analysis of *IFNB1* and *IFNL1* in AAVS1 or MAVS KO HAE-ALI cultures infected with IAV (diluted 1:20 in DMEM) or left uninfected for 16 hours. Each data point represents an independent culture (n=3) derived from the same donor. For *IFNL1*, some uninfected cultures had undetectable levels. **B)** RT-qPCR analysis on ISGs in cultures infected with IAV (diluted 1:20 in DMEM) or left uninfected for 16 hours. Each data point represents an independent culture (n=3) derived from the same donor. **C)** Western blot analyzing protein levels of MAVS, various ISGs and viral (NS1) protein in AAVS1 or MAVS KO HAE-ALI cultures 48 h post-infection with IAV. Vinculin (VCL) was used as a loading control. Each lane represents an independent culture (uninfected n=1, infected n=3) derived from the same donor. **D)** hCXCL-10 ELISA performed on the apical mucus wash from AAVS1 or MAVS KO HAE-ALI cultures 48 hours after infection with IAV (diluted 1:20 in DMEM) or left uninfected. Each data point represents an independent culture (uninfected n=1, infected n=6) derived from the same donor. **E)** Basolateral media underneath the transwell membrane from the same experiment as in (D) was also analyzed using a hCXCL-10 ELISA. Each data point represents an independent culture (uninfected n=1, infected n=6) derived from the same donor. **F)** RT-qPCR analysis of viral RNA from three different viral segments from AAVS1 or MAVS KO HAE-ALI cultures infected with IAV (diluted 1:20 in DMEM) for 48 hours. Statistical difference was determined by an unpaired Welch’s t-test; “ns” indicates p-value > 0,05. **G)** Western blot analysis of MAVS and viral (NS1) protein levels in AAVS1 or MAVS KO HAE-ALI cultures infected with IAV (diluted 1:20 in DMEM) or SeV (30 HAU/mL) or left uninfected for 16 hours. VCL was used as a loading control. Each lane represents an independent culture (n=3) derived from the same donor. Bars represent mean ± s.e.m. Unless otherwise stated, statistical differences were determined by multiple unpaired t-tests with FDR correction using the two-stage step-up method of Benjamini, Krieger and Yekuteili (desired FDR = 5%). Q-values are shown above each comparison. Statistical analysis was not performed for uninfected groups (A-B and D-E).

